# Discrete viscoelastic model of deformable immune cell migration through confined tumor microchannels

**DOI:** 10.64898/2025.12.22.695944

**Authors:** David Garcia-Navarro, Jack Zhang-Zhou, Jose Manuel Garcia-Aznar, Daniel Camacho-Gomez

**Affiliations:** Department of Mechanical Engineering, Multiscale in Mechanical and Biological Engineering (M2BE), Aragon Institute of Engineering Research (I3A), University of Zaragoza, Zaragoza, Spain; Integrated Mathematical Oncology Department, Moffitt Cancer Center, Tampa, FL

**Keywords:** cell deformation, particle-based simulations, mechanical evaluation, viscoelastic models

## Abstract

Understanding the mechanical behavior of immune cells as they migrate through the complex environment of solid tumors may be crucial for improving immunotherapy strategies. In this work, we present a particle-based model that captures the large deformations of single immune cells migrating through confined microchannels, which mimic the narrow tumor micropores. In our model, the nucleus, cytoplasm, and membrane are simulated through discrete particles with varying interacting forces that account for the differences in the mechanical properties of the discrete parts of the cell. We compare and validate the model with experimental measurements from microfluidic devices, providing relevant mechanistic conclusions into the effects of changes in cellular stiffness on migration efficiency.

## 1. Introduction

The burden of cancer represents an increasing global challenge, accounting for 16.8% of all deaths worldwide with an increasing incidence [1, 2]. Innovative approaches are being developed to improve patient survival rates. Among these promising treatments, immunotherapy has shown success by enhancing the immune system’s ability to recognize and attack malignant cells [3, 4]. These therapies have already improved survival outcomes in cancers with very unfavorable prognoses, also increasing the number of patients who could benefit from these treatments [5]. One of these therapies is based on the genetic modification of T lymphocytes to express a

Chimeric Antigen Receptor (CAR), which enables these cells to recognize and attack tumor cells more effectively [6–13]. Currently, this CAR-T therapy has proven efficacy in cases such as B-cell leukemia or lymphomas, but not in solid tumors or other hematological malignancies [14]. This limited effectiveness in solid tumors might be attributed to several factors, including the lack of a universal tumor-specific antigen, the immunosuppressive environment within the tumor, T-cell exhaustion, or the lack of immune CAR-T cell infiltration into the tumor core, hindered by both physical and biological barriers [4, 5, 14–16]. Although clinical trials remain the only definitive way to evaluate the efficacy and safety of immunotherapies, identifying the mechanical and biophysical factors that hinder immune cell infiltration into solid tumors is essential to understanding their limited effectiveness.

A way to study the infiltration of immune cells into the tumor is through *in vitro* microfluidic experiments [17–20], which enable precise control of the microenvironment to assess how its mechanical properties affect cell deformability, migration, and internal organization. Using this approach, Zhang-Zhou et al. study how immune cells migrate through narrow microchannels, which mimic the penetration of these cells through tumor micropores. Their results suggest that the expression of CARs may increase cell stiffness, triggered by cellular remodeling resulting from T-cell genetic modification [21, 22], which might be the reason behind the poor infiltration of immune cells through tumor micropores.

In this context, computational models can provide powerful tools to investigate these phenomena, enabling the characterization of the mechanical properties of the cell and its microenvironment [23], providing insights that are often inaccessible through experiments alone. Computational models have been developed to study cell migration, including continuous models [24–27], Agent-Based Models (ABM) [28, 29], also combined with deep reinforcement learning [30], and other hybrid approaches [23, 31, 32]. For instance, Jamali et al. [33] develop a Subcellular Element Model (SEM) to study how cell deformation influences the global morphology of epithelial monolayers. Escribano et al. [34] propose a model focused on collective cell migration regulated by substrate stiffness, i.e., durotaxis. More recently, Chattaraj et al. [35] design a framework to analyze cell deformation in situations with *in vivo* and *in vitro* applications, such as morphogenesis. Other models, such as the one developed by Alawadhi et al. [36], simulate the amoeboid migration of T cells using a bead-based model that accounts for the hydraulic nuclear piston mechanism and the role of the uropod as a stabilizing element during migration in an ECM with high obstacle density. However, these models do not account for the entire cell deformation, or they do not incorporate the cell membrane, despite increasing evidence that membrane mechanics and dynamics play a critical role in regulating migratory behavior [37–39]

Here, we present an SEM to simulate the migration of single immune cells through microchannels of different sizes, which represent tumor micropores, accounting for large cell deformation. To achieve this, the model considers not only the three main parts of the cell (nucleus, cytoplasm, and membrane) but also the interaction with the walls of the microchannel. Thus, interactions between particles determine the global viscoelastic behavior of the cell, enabling it to deform and penetrate into the narrow microchannels. We apply this model to simulate the migration of single

T-cells and CAR-T cells in confined spaces as developed by Zhang-Zhou et al. [17]. We calibrate and validate the model by reproducing cell deformation and velocity during migration with the available experimental measurements. This allows us to evaluate the mechanical behavior of different cell types, which in turn allows us to investigate how the generation of T cells into CAR-T cells affects their mechanical properties and their ability to infiltrate into tumors.

In this paper, we first introduce the computational model employed, highlighting its main features and the specific cellular behavior it aims to reproduce. We then describe the mechanical interactions and driving forces governing cell dynamics, followed by the experimental procedures that provide the data for model calibration and validation. Finally, we present and discuss the main simulation results, addressing the model’s limitations and potential applications.

## 2. Methods

### 2.1. Computational framework

We present a particle-based model to simulate the migration of immune cells through confined microchannels (Fig. 1**a**). This model consists of an Off-Lattice SEM model with a continuous description of cell position, where each cell is discretized by multiple interacting particles.

**Fig. 1.**
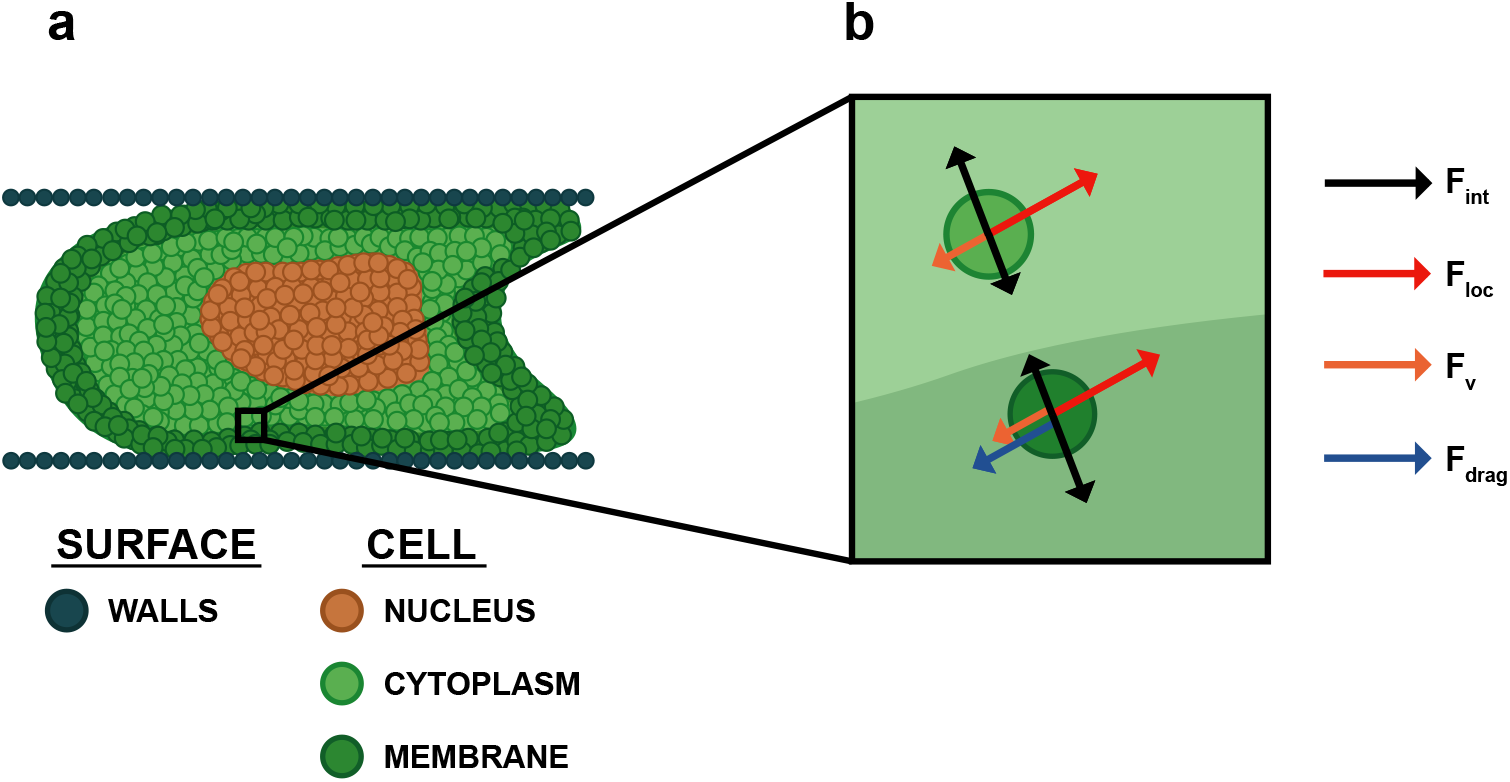
Model overview. a. Discrete approximation of the cell, including three different types of particles representing the main parts of the cell (nucleus, cytoplasm, and membrane), and an additional particle type which discretizes the wall surface. b. Schematic representation of the main forces in the model. *F*_*int*_ denotes the interaction force between particles, which can be adhesive or repulsive depending on their relative Euclidean distance, *F*_*loc*_ represents the active force driving cell migration, *F*_*v*_ accounts for the viscous behavior of the cell arising from internal resistance to relative motion between its components, and *F*_*drag*_ denotes the viscous drag force due to friction of the cell with the surrounding viscous fluid.

The model represents the viscoelastic behavior of the cell, incorporating elastic components to reproduce the cell’s resistance to deformation and a viscous component to allow a progressive deformation of the cell to enter into the tumor ducts and flow through them, modifying and redistributing the inside of the cells.

For this purpose, we define four types of particles. The first represents the substrates over which the cell migrates; these have fixed positions and very stiff bonds to avoid deformation. The second one models the cell nucleus, also connected by stiff joints, but allowing mobility. The third discretizes the cytoplasm, characterized by significantly weaker joins, representing its inherent fluidity. The last type corresponds to membrane particles, which show stiff in-plane connections due to the membrane’s high extension strength, but weaker interactions with cytoplasmic particles, reflecting its low shear strength. This type ultimately determines the overall deformation of the cell.

We integrate our numerical approach into *PhysiCell* [40], an open-source framework for simulating 3D multicellular systems through ABMs, to enhance its capabilities by enabling the simulation of cell deformation, where each cell is discretized into multiple interacting particles of different types that represent its main structural components. To ensure biologically realistic behavior, we develop a parameter-calibration method that reproduces the mechanical responses of the nucleus, cytoplasm, and membrane, while also enforcing numerical constraints that prevent non-physical configurations that would otherwise lead to cell collapse or death.

### 2.2. Mechanical interactions

To study cell deformation, we describe the temporal evolution of each particle that discretizes the cell. Let *δ*_*i*_ = {*N, M, C, S*}, being the *i* particle of the subsets *δ*, which can be nucleus (N), membrane (M), cytoplasm (C), or surface (S), the position of 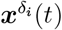 is calculated from the force balance:

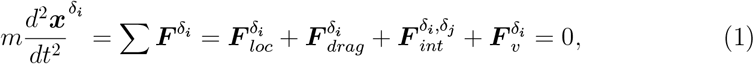

where 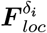 represents the locomotive force, 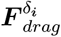 is the friction with the ECM, 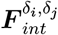 is the elastic interaction between the particles *δ*_*i*_ and *δ*_*j*_ and 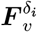 corresponds to the viscous interaction of the particle with the inside of the cell (Fig. 1**b**).

The locomotive force is the active force exerted by the cell that regulates its movement, so it is applied to all the particles. Hence, *δ* can take any of the values of the subset {*N, M, C*.}

Drag force results from the friction of the cell in contact with a viscous fluid, so it only applies to membrane particles *δ* = {*M}*. We calculate it according to the Stokes law:

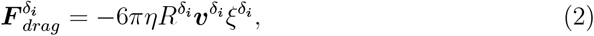

where *η* is the dynamic viscosity of the ECM, 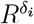 and 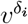 corresponds to the radius and velocity of the particle respectively, and 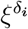 equals 1 if it is a membrane particle and 0 otherwise.

Interaction forces arise between nearby particles to simulate cell stiffness, i.e., its opposition to deformation. To calculate this force, we define a maximum adhesion distance (*R*_*A*_), which determines the maximum distance considered between neigh-boring cells. This interacting force can be adhesive or repulsive, depending on the distance between particles 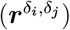.

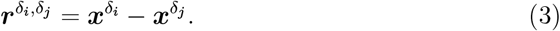

As this force acts on all cell particles and also accounts for cell–surface interactions, *δ*_*i*_ can take any value from the subset {*N, M, C*}, while *δ*_*j*_ can take values from the subset {*N, M, C, S*}.

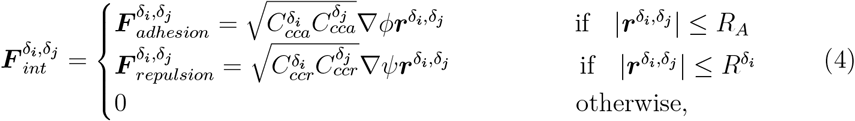

where 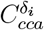 and 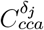 correspond to the adhesion coefficients of the particles *δ*_*i*_ and 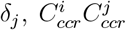 are the repulsion coefficients of the particles, ∇*ϕ* and ∇*ψ* correspond to the potential law that regulates the adhesion and repulsion between particles, respectively, as defined in Ghaffarizadeh et al..

Viscous force introduces the viscous behavior of the cell, that is, the internal resistance exerted by the cell to the relative movement of its parts, including the different organelles and cytosol that are not explicitly modeled.

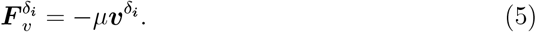

So, it is proportional to the internal viscosity of the cell *μ* and the velocity of the particle 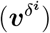. Since this force is applied to the entire cell, *δ* can take any of the values of the subset {*N, M, C*}.

### 2.3. Experimental Setup

To recreate the mechanical barriers present in the solid tumor stroma, a novel PDMS-based microfluidic device is designed, as shown by Zhang-Zhou et al. The device geometry is optimized through microfluidic simulations in COMSOL Multi-physics®, and consists of two main channels interconnected by microchannels of 2, 4, 6, and 8 μm in width, in quantities of 37, 95, 75, and 28, respectively, to mimic the structural heterogeneity of solid tumors. The height and length of all microchannels are fixed at 6 and 200 μm, respectively (Fig. 2**a**). These microchannel devices are coated with collagen type I (20 μg*/*μL) and ICAM-I (10 μg*/*μLL) for 1 h at 37^°^ to enhance cell adhesion. In this experiment, T cells and CAR-T cells—isolated from PBMCs obtained via Ficoll-Paque gradient centrifugation from a healthy donor’s blood supplied by the Blood and Tissue Bank of Aragon—were seeded into one of the main channels at a concentration of 2 x 10^6^ cells/mL. Cells were allowed to migrate through the perpendicular microchannels toward the opposite main channel. Cell migration was monitored using time-lapse live-cell microscopy (Axio Observer 7, Zeiss, Germany; Eclipse Ti-E inverted microscope, Nikon, The Netherlands).

**Fig. 2.**
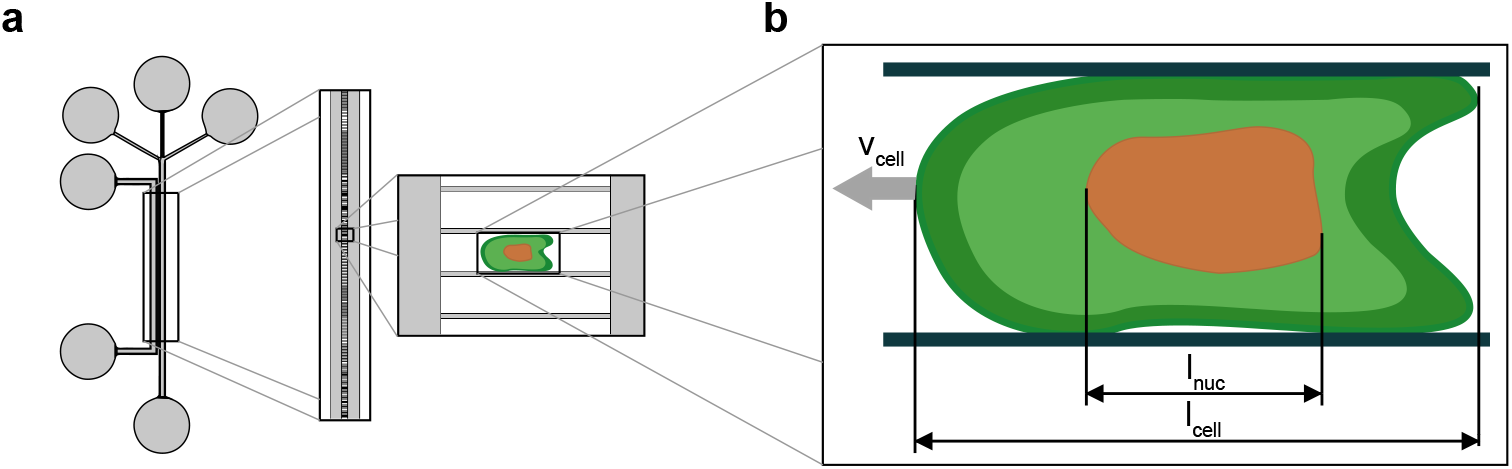
Experimental setup. a. PDMS-based microfluidic device consisting of two main channels interconnected by microchannels of 2, 4, 6, and 8 μm in width. b. Experimental measurements used to calibrate and validate the model. *v*_*cell*_ represents the cell migration velocity inside the microchannel, whereas *l*_*cell*_ and *l*_*nuc*_ denote the cell and nucleus lengths during migration.

Brightfield images were captured via a 10× objective at 50–70 s intervals for 12 hours. These recordings provided quantitative information on cell velocity and morphology during confined migration, including the length of both the nucleus and the whole cell (Fig. 2**b**). Additionally, cellular energy consumption was analyzed by labeling immune cell mitochondria with MitoTracker™Red CMXRos (Thermo Fisher Scientific), following the manufacturer’s instructions.

### 2.4. Model calibration with experimental measurements

We calibrate our model by reproducing cell deformation and velocity during confined migration. First, we define the specific discrepancies (*x*_*i*_) between experimental measurements [2**b**] and computational predictions. These deviations focus on cell morphology, migration dynamics, and numerical stability:

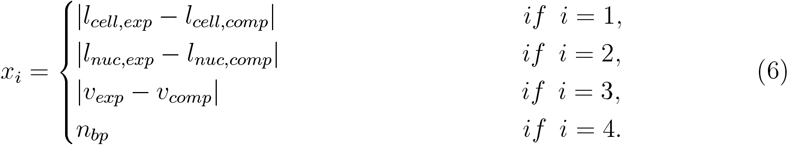

In our computational model, *l*_*cell,comp*_ and *l*_*nuc,comp*_ are calculated as the Euclidean distance between the two most distant particles of the cell and the nucleus, respectively. The velocity *v*_*comp*_ is computed as the mean velocity of all cell particles after passing the curved inlet of the microchannel. Additionally, we track *n*_*bp*_, representing the number of particles detached from the main cell body, which is penalized in the objective function. To translate these raw deviations into a normalized metric that accounts for the intrinsic biological variability of the experiments (n=4), we employ a hyperbolic tangent error function:

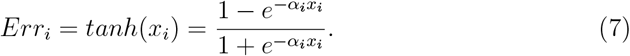

Here, the scaling factors *α*_*i*_ are calibrated to ensure the function saturates at a target threshold. This formulation is strategically chosen to strongly penalize significant discrepancies while dampening the influence of minor deviations arising from experimental noise, preventing excessive penalties for acceptable biological variance. The global objective function is subsequently defined as the arithmetic mean of these four normalized error components:

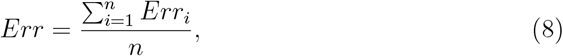

where the individual error terms correspond to the cell length, nucleus length, cell velocity, and detached particles errors, respectively. Minimizing this total error function allows us to identify the optimal set of parameters that define the cell’s mechanical interactions and deformation. These parameters include the internal cell viscosity (*μ*), the locomotive force driving migration (*F*_*Loc*_), and the discrete adhesion and repulsion coefficients (*K*^*δ*^, where *α* denotes adhesion or repulsion, and *δ* specifies the particle type: nucleus, membrane, or cytoplasm). Finally, we perform a hyperparameter optimization with the open-source framework Optuna [41]. Specifically, we employ Bayesian optimization, using the Tree-structured Parzen Estimator (TPE) sampler, which is particularly effective for large parameter spaces and computationally expensive model evaluations [42–45].

## 3. Results

### 3.1. Model calibration and validation

We calibrate the model with experimental data to evaluate cell mechanics. For this purpose, we develop a robust method that validates the results obtained to minimize an error function that includes the three main metrics (migration speed, cell length, and nucleus length) that we aim to reproduce, as these are the ones that allow us to adjust the dynamics and morphology of migration. As shown in Fig. 3**a**, we optimize the model with an error below 5% for all experimental conditions, indicating that the model achieves a sufficiently accurate fit. In addition, the parity plots in Fig. 3**b-d** show that the computationally obtained values closely match the experimental measurements, with most points lying near the 45° identity line. The largest deviation is observed for the nucleus length (Fig. 3**d**), where the model slightly underestimates the experimental values, although the discrepancy remains within acceptable limits. No systematic overestimation or underestimation is observed for the other metrics. Overall, these results indicate that the model reliably and accurately reproduces the essential features of confined cell migration.

**Fig. 3.**
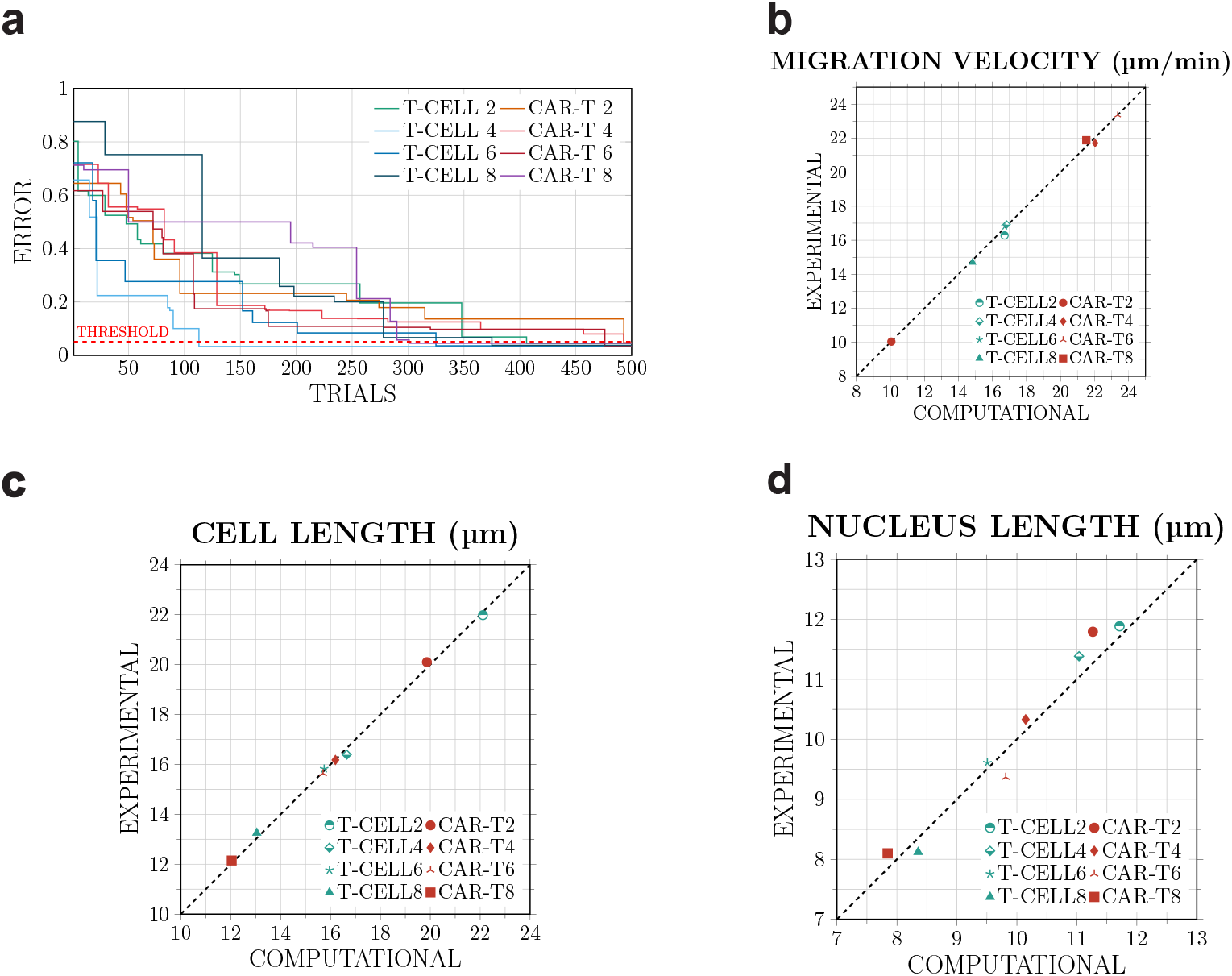
Validation of the model. a. Evolution of the error function against the trials for the different simulations. b. Parity plot, which compares the computational predictions with the experimental measurements by plotting one against the other, of cell migration velocity. c. Parity plot of cell length. d. Parity plot of nucleus length.

### 3.2. Predictive mechanical evaluation of T-Cells

To mechanically assess T-cells, we use the optimization method described above to obtain the set of parameters that best reproduces the experimental observations. Specifically, we focus on the adhesion and repulsion constants of the nucleus, cytoplasm, and membrane; the locomotion force required for migration through the microchannels; and the internal cell viscosity. We calibrated the model parameters and obtained an estimation of the adhesion and repulsion constants (Fig. 4**a-b**), which vary depending on the degree of confinement. For both types of constants, the highest values are generally observed in the intermediate channel widths (6 and 8 μm). In addition, the repulsion constants are considerably higher than the adhesion constants, in some cases by up to two orders of magnitude. Fig. 4**c** shows the locomotion force obtained for each confinement condition. The largest forces correspond to the least confined situations (5.35e-03, 5.89e-03, and 5.29e-03 μN for the 4, 6, and 8 μm channels, respectively) whereas the smallest force corresponds to the 2 μm channel (3.52e-03 μN). Similarly, the viscosity values displayed in Fig. 4**d** are highest for the intermediate channel widths, reaching values approximately two to three times larger than those obtained for the most confined and least confined conditions. Finally, Fig. 4**e** shows that the morphology adopted by the cell during migration through the 4 μm microchannel in the simulations (right) closely matches that observed experimentally (left), indicating that the model reproduces the characteristic deformation patterns associated with confined migration in the microchannels. Therefore, the model allows us to mechanically evaluate cells during their confined migration and compare the behavior of different types of cells.

**Fig. 4.**
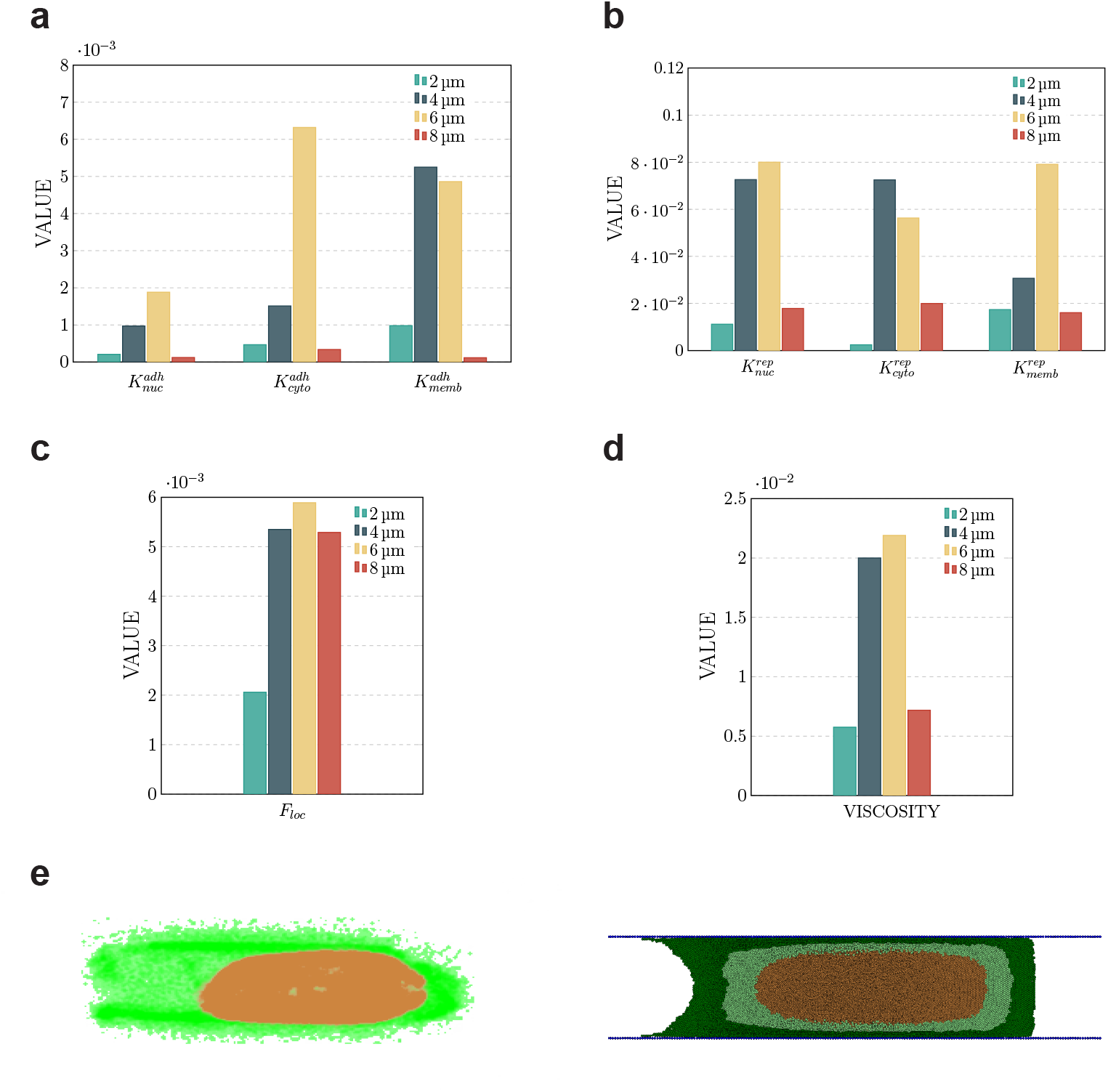
Mechanical evaluation of T-cells. a. Obtained values for adhesion (*k*_*adh*_) constants for the nucleus (N), cytoplasm (C), and membrane (M) in the different microchannels. b. Obtained values for repulsion (*k*_*rep*_) constants for N, C and M in the different microchannels. c. Locomotive force in the studied situations. d. Internal cell viscosity in the different microchannels. e. Representation of cell deformation in the 4 μm microchannel, obtained experimentally (left) and computationally (right).

### 3.3. Predictive mechanical evaluation of CAR-T cells

Additionally, to ensure that the model can simulate different types of cells and to be able to make comparisons between them, in this subsection, we mechanically evaluate CAR-T cells. Fig. 5**a-b** shows that the value of the adhesion and repulsion constants varies with respect to the different tube sizes, and that the repulsion constants are significantly higher than the adhesion constants. For CAR-T cells, the highest repulsion constants are observed in the 2 μm microchannels. Similarly, the locomotion force (Fig. 5**c**) and internal viscosity (Fig. 5**d**) reach their maximum values under the same confinement conditions Fig. 5**e** presents the comparison of the geometries adopted by the cell in the experiments (left) and in the simulations (right) in the 4 μm microchannel. These results confirm that the model can reproduce the mechanical behavior of different cell types under confinement, providing a foundation for the comparative analysis between T cells and CAR-T cells presented in the following subsection.

**Fig. 5.**
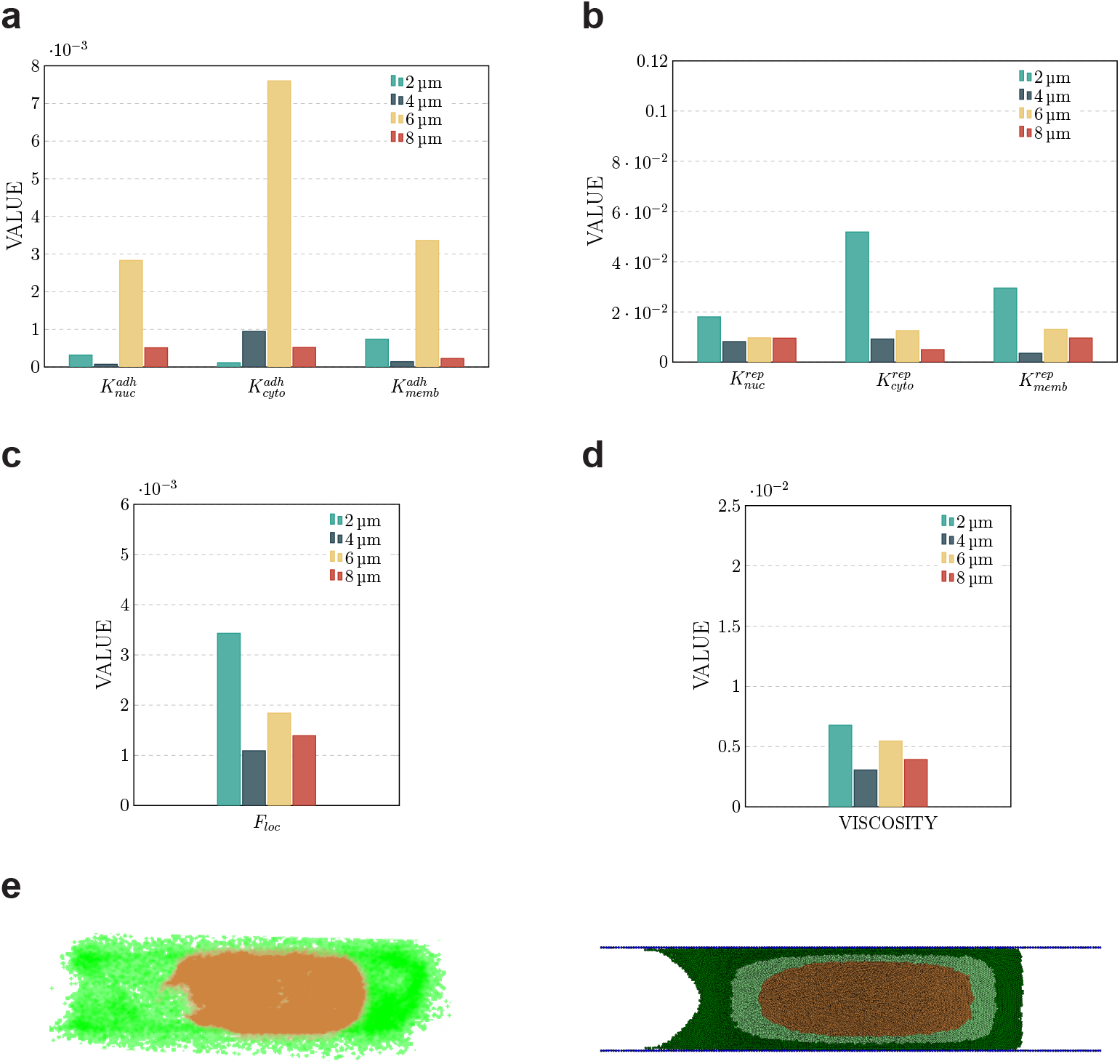
Mechanical characterization of CAR-T cells. a. Obtained values for adhesion (*k*_*adh*_) constants for the nucleus (N), cytoplasm (C), and membrane (M) in the different microchannels. b. Obtained values for repulsion (*k*_*rep*_) constants for N, C and M in the different microchannels. c. Locomotive force in the studied situations. d. Internal cell viscosity in the different microchannels. e. Representation of cell deformation in the 4 μm microchannel, obtained experimentally (left) and computationally (right).

### 3.4. A comparative analysis between T and CAR-T cells

Finally, we compare the values of the model parameters that show significant differences between two types of immune cells used in cancer treatments. As illustrated in Fig. 6**a-e**, for simulations in the intermediate and low confinement conditions (4, 6, and 8 μm microchannels), all parameters (adhesion and repulsion constants, locomotion force, and intracellular viscosity) are higher for T cells than for CAR-T cells. In contrast, in the highest confinement condition (2 μm microchannel), the parameter values are higher for CAR-T cells. Fig. 6**f** presents the distribution of experimentally measured migration velocities for T cells and CAR-T cells ([17]). T cells exhibit relatively constant velocities across the different channel widths: 13.38, 17.15, 17.85, and 14.76 μm*/*min for the 2, 4, 6, and 8 μm tubes, respectively. In contrast, CAR-T cells show a reduced velocity in the 2 μm microchannel (9.89 μm*/*min) and higher velocities in the 4, 6, and 8 μm channels (21.75, 23.15, and 21.94 μm*/*min, respectively). These results clearly indicate that the mechanical parameters differ between T cells and CAR-T cells under the same confinement conditions, and that these differences are experimentally reflected in the observed behavior.

**Fig. 6.**
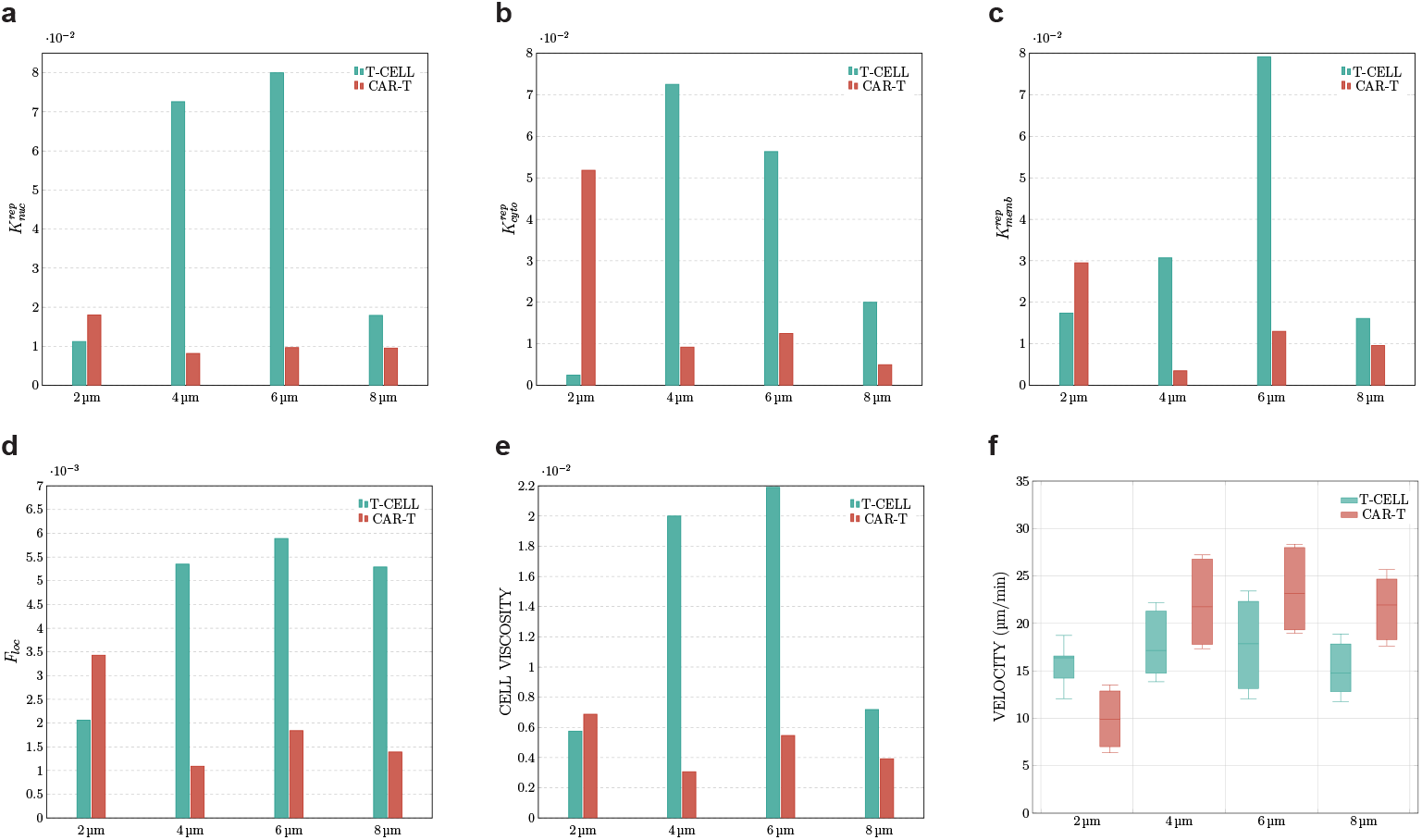
Comparison of the properties with major differences between T-cells and CAR-T cells. a. Comparison of the repulsion constants in nucleus particles 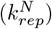. b. Comparison of the repulsion constants in cytoplasm particles 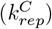. c. Comparison of the repulsion constants in membrane particles 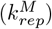. d. Comparison of cell locomotion forces. e. Comparison of cell internal viscosities. f. Representation of the distribution of the migration velocities of the two cell types in the different microchannels.

## 4. Discussion

In this paper, we present a computational approach to study the role that cell mechanics play during confined migration and understand how different immune cells infiltrate through narrow tumor micropores. Unlike other continuous or discrete models that often treat cells as rigid or uniform entities, our off-lattice SEM explicitly discretizes the cell into a large number of particles. This approach allows us to reproduce the large deformations required for the cell to enter the microchannel. The model considers three different types of interacting particles representing the main cellular components (nucleus, cytoplasm, and membrane), as well as an additional type used to model the microchannel surface. This combination of particles enables an accurate representation of cellular viscoelasticity. The model is calibrated and validated by reproducing the morphology and dynamics of single immune cell migration through 2, 4, 6 and 8 μm microchannels, as developed by Zhang-Zhou et al. (Fig. 2**b**), obtaining a high degree of accuracy, with prediction errors consistently below 5% (Fig. 3**a**). This reliability establishes the model not merely as a simulation tool, but as a predictive platform for evaluating cellular mechanics. With this model we characterize different cell types, such as T cells (Fig. 4) or CAR-T cells (Fig. 5) during migration. In this evaluation of mechanical properties, we can observe differences depending on the level of confinement, determined by the size of the microchannel through which the cell must migrate. This has been studied previously [46–48], which relate changes in cellular structure and function caused by confinement. Other studies also show lower cell stiffness in more confined environments [49, 50]. We can observe this phenomenon in the evolution of the repulsion constants of the different parts of the cell (since greater repulsive force entails a lower capacity for cell deformation, as it hinders movements between particles), which generally increase as the microchannel size increases from 2 to 6 μm. However, in the 8 μm tube, given the initial size of the cells (10.61 μm), confinement is much lower, so it may not fit this pattern.

Therefore, we consider the most important comparison to be of mechanical properties under the same confinement conditions. We can observe for the majority of the simulations (migration through 4, 6 and 8 μm microchannels) that the repulsion constants for all parts of the cell considered are greater in the case of the T cells, whereas in the 2 μm microchannel we can see a clear change of this trend, since the repulsion constants are larger for the CAR-T cells, which would indicate a greater stiffness of the CAR-T cell (Fig. 6**a-c**). Also, looking at the comparison of locomotion forces (Fig. 6**d**), while in the 4, 6 and 8 μm cases they are larger for the T cells, in the 2 μm microchannel they are higher for the CAR-T cells. We hypothesize that this is because, as the cell is stiffer, the normal force it exerts on the surface of the microchannel is larger, which produces a greater frictional force, and, consequently, a larger locomotive force must be applied. Finally, cellular viscosity follows the same trend (Fig.6**e**), which would indicate a greater importance of the viscous component of behavior for CAR-T cells under extreme confinement.

From the experimental data [17], we can observe both behaviors. First, migration velocities follow a similar pattern to our simulations: in 4, 6, and 8 μm microchannels, CAR-T cells migrate faster, whereas in the 2 μm channel, T cells exhibit higher velocities (Fig. 6**f**). Second, CAR-T cells show increased mitochondrial activity during migration through the 2 μm channel, indicating a higher energetic demand. Thus, both trends support our hypothesis that, under extreme confinement, CAR-T cells exhibit greater stiffness than T cells.

However, we have to keep in mind that several simplifications were introduced to reduce the complexity of the model while preserving its validity. First, the cell is discretized into its three main components (nucleus, cytoplasm, and membrane), each treated as a homogeneous medium, without explicitly representing organelles or other subcellular structures. Although simplified, this approach captures the overall dynamics of cell deformation and migration. Second, we assume that the mechanical and behavioral properties of the cell remain constant throughout migration, despite potential changes during migration. In addition, all simulated cells were the same size, despite the significant biological variability that exists. Future work will incorporate different cell geometries sampled from a statistically representative distribution to better reflect this heterogeneity. While the variables of the model provide a qualitative insight into the corresponding cellular behavior, performing an AFM assay to measure the actual mechanical properties of the cells will allow us to establish direct correlations with the variables in the model. It would also be of interest to integrate this cell deformation model into geometries that mimic the morphology of solid tumors and incorporate biochemical cues known to regulate immune cell infiltration.

Overall, the developed model shows strong potential for studying cell deformation during confined migration. Thanks to this evaluation of cell mechanical deformation, we have seen not only the different cell stiffness depending on the degree of confinement for both types of cells, but also that CAR-T cells are stiffer than T-cells for the narrowest microchannels (2 μm) and less stiffer for the wider microchannels, which would be supported by the literature [49, 50]. Finally, we have discovered a direct relationship between cell stiffness and lower migration speeds, which may be explained by the greater pressure exerted by the cell on the microchannel, thus increasing friction with its surface.

## 5. Aknowledgements

This paper is part of a project that has received funding from the European Research Council (ERC) under the European Union’s Horizon 2020 research and innovation programme (ICoMICS grant agreement No 101018587)”.

## 6. Author contributions

Conceptualization: D. G-N., D. C-G., and JM. G-A.; Computational design: D. G-N. and D. C-G.; Computational implementation: D. G-N; Experimental implementation: J. Z-Z.; Funding acquisition: JM.G-A.; Original draft: D. G-N.; Review and editing: D. G-N., D. C-G., J. Z-Z., and JM. G-A.

## References

[1] Freddie Bray, Mathieu Laversanne, Hyuna Sung, Jacques Ferlay, Rebecca L. Siegel, Isabelle Soerjomataram, and Ahmedin Jemal. Global cancer statistics 2022: Globocan estimates of incidence and mortality worldwide for 36 cancers in 185 countries. CA: A Cancer Journal for Clinicians, 74:229–263, 5 2024. ISSN 0007-9235. doi: 10.3322/CAAC.21834. URL https://doi.org/10.3322/CAAC.21834.

[2] Isabelle Soerjomataram and Freddie Bray. Planning for tomorrow: global cancer incidence and the role of prevention 2020–2070. Nature Reviews Clinical Oncology, 18:663–672, 10 2021. ISSN 17594782. doi: 10.1038/s41571-021-00514-z. URL https://doi.org/10.1038/s41571-021-00514-z.

[3] Shuzhen Tan, Dongpei Li, and Xiao Zhu. Cancer immunotherapy: Pros, cons and beyond. Biomedicine & Pharmacotherapy, 124:109821, 4 2020. ISSN 0753-3322. doi: 10.1016/J.BIOPHA.2020.109821. URL https://doi.org/10.1016/J.BIOPHA.2020.109821.

[4] Jaechang Kim, Ruby Maharjan, and Jonghyuck Park. Current trends and innovative approaches in cancer immunotherapy. AAPS PharmSciTech 2024 25:6, 25:1–17, 7 2024. ISSN 1530-9932. doi: 10.1208/S12249-024-02883-X. URL https://doi.org/10.1208/S12249-024-02883-X.

[5] Alex D. Waldman, Jill M. Fritz, and Michael J. Lenardo. A guide to cancer immunotherapy: from t cell basic science to clinical practice. Nature Reviews Immunology, 20:651–668, 11 2020. ISSN 14741741. doi: 10.1038/s41577-020-0306-5. URL https://doi.org/10.1038/s41577-020-0306-5.

[6] Qingyang Zhang, Jieming Ping, Zirui Huang, Xiaoli Zhang, Jingyi Zhou, Gangyang Wang, Shaoyang Liu, and Jianjun Ma. Car-t cell therapy in cancer: Tribulations and road ahead. Journal of Immunology Research, 2020:1924379, 1 2020. ISSN 2314-7156. doi: 10.1155/2020/1924379. URL https://doi.org/10.1155/2020/1924379.

[7] Lihong Wang, Lufang Zhang, Louisa Chard Dunmall, Yang Yang Wang, Zaiwen Fan, Zhenguo Cheng, and Yaohe Wang. The dilemmas and possible solutions for car-t cell therapy application in solid tumors. Cancer Letters, 591:216871, 6 2024. ISSN 0304-3835. doi: 10.1016/J.CANLET.2024.216871. URL https://doi.org/10.1016/J.CANLET.2024.216871.

[8] Sai Rohit Reddy, Adiona Llukmani, Ayat Hashim, Dana R. Haddad, Dutt S. Patel, Farrukh Ahmad, Majdi Abu Sneineh, Domonick K. Gordon, Sai Rohit R. Mekala, Adiona Llukmani, Ayat Hashim, Dana R. Haddad, Dutt S. Patel, Farrukh Ahmad, Majdi Abu Sneineh, and Domonick K. Gordon. The role of chimeric antigen receptor-t cell therapy in the treatment of hematological malignancies: Advantages, trials, and tribulations, and the road ahead. Cureus, 13, 2 2021. doi: 10.7759/CUREUS.13552. URL https://doi.org/10.7759/CUREUS.13552.

[9] Xiaojuan Shi, Daiqun Zhang, Feng Li, Zhen Zhang, Shumin Wang, Yujing Xuan, Yu Ping, and Yi Zhang. Targeting glycosylation of pd-1 to enhance car-t cell cytotoxicity. Journal of Hematology and Oncology, 12:1–4, 11 2019. ISSN 17568722. doi: 10.1186/S13045-019-0831-5. URL https://doi.org/10.1186/S13045-019-0831-5.

[10] Sangita Dey, Moodu Devender, Swati Rani, and Rajan Kumar Pandey. Recent advances in car t-cell engineering using synthetic biology: Paving the way for next-generation cancer treatment. Advances in Protein Chemistry and Structural Biology, 140:91–156, 1 2024. ISSN 1876-1623. doi: 10.1016/BS.APCSB.2024.02.003. URL https://doi.org/10.1016/BS.APCSB.2024.02.003.

[11] Ruihao Huang, Xiaoping Li, Yundi He, Wen Zhu, Lei Gao, Yao Liu, Li Gao, Qin Wen, Jiang F. Zhong, Cheng Zhang, and Xi Zhang. Recent advances in car-t cell engineering. Journal of Hematology and Oncology, 13:1–19, 7 2020. ISSN 17568722. doi: 10.1186/S13045-020-00910-5. URL https://doi.org/10.1186/S13045-020-00910-5.

[12] Kusala Anupindi, Julia Malachowski, Isabella Hodson, Daniel Zhu, Carl H. June, and Bruce L. Levine. The next innovations in chimeric antigen receptor t cell immunotherapies for cancer. Cytotherapy, 5 2025. ISSN 1465-3249. doi: 10.1016/J.JCYT.2025.05.010. URL https://doi.org/10.1016/J.JCYT.2025.05.010.

[13] Elmar Jaeckel, Scott L. Friedman, Michael Hudecek, and Ulrike Protzer. Chimeric antigen receptor (car) t-cell therapy: Engineering immune cells to treat liver diseases. Journal of Hepatology, 6 2025. ISSN 0168-8278. doi: 10.1016/J.JHEP.2025.06.007. URL https://doi.org/10.1016/J.JHEP.2025.06.007.

[14] Robert C Sterner and Rosalie M Sterner. Car-t cell therapy: current limitations and potential strategies. Sterner and Sterner Blood Cancer Journal, 11:69, 2021. doi: 10.1038/s41408-021-00459-7. URL https://doi.org/10.1038/s41408-021-00459-7.

[15] Lydia G. White, Hannah E. Goy, Alinor J. Rose, and Alexander D. McLellan. Controlling cell trafficking: Addressing failures in car t and nk cell therapy of solid tumours. Cancers, 14, 2 2022. ISSN 20726694. doi: 10.3390/CANCERS14040978. URL https://doi.org/10.3390/CANCERS14040978.

[16] Andrea Ladányi. Prognostic and predictive significance of immune cells infiltrating cutaneous melanoma. Pigment Cell and Melanoma Research, 28:490–500, 9 2015. ISSN 1755148X. doi: 10.1111/PCMR.12371. URL https://doi.org/10.1111/PCMR.12371.

[17] Jack Zhang-Zhou, Nieves Movilla Meno, Carmen Oñate Salafranca, Maria Jose Gomez-Benito, Pedro Enrique Guerrero, Julian Pardo Jimeno, and Jose Manuel García-Aznar. Car-t cells are more affected than t lymphocytes by mechanical constraints: A microfluidic-based approach. Life Sciences, 363:123335, 2025. doi: 10.1016/j.lfs.2024.123335. URL https://doi.org/10.1016/j.lfs.2024.123335.

[18] Colin D. Paul, Daniel J. Shea, Megan R. Mahoney, Andreas Chai, Victoria Laney, Wei Chien Hung, and Konstantinos Konstantopoulos. Interplay of the physical microenvironment, contact guidance, and intracellular signaling in cell decision making. FASEB Journal, 30:2161–2170, 6 2016. ISSN 15306860. doi: 10.1096/FJ.201500199R. URL https://doi.org/10.1096/FJ.201500199R.

[19] Chieh Ren Hsia, Jawuanna McAllister, Ovais Hasan, Julius Judd, Seoyeon Lee, Richa Agrawal, Chao Yuan Chang, Paul Soloway, and Jan Lammerding. Confined migration induces heterochromatin formation and alters chromatin accessibility. iScience, 25:104978, 9 2022. ISSN 2589-0042. doi: 10.1016/J.ISCI.2022.104978. URL https://doi.org/10.1016/J.ISCI.2022.104978.

[20] Eric M. Balzer, Ziqiu Tong, Colin D. Paul, Wei Chien Hung, Kimberly M. Stroka, Amanda E. Boggs, Stuart S. Martin, and Konstantinos Konstantopoulos. Physical confinement alters tumor cell adhesion and migration phenotypes. FASEB Journal, 26:4045–4056, 10 2012. ISSN 15306860. doi: 10.1096/FJ.12-211441. URL https://doi.org/10.1096/FJ.12-211441.

[21] Rui Li, Chao Ma, Haogang Cai, and Weiqiang Chen. The car t-cell mechanoimmunology at a glance. Advanced Science, 7:2002628, 12 2020. ISSN 21983844. doi: 10.1002/ADVS.202002628. URL https://doi.org/10.1002/ADVS.202002628.

[22] Yiwei Xiong, Kendra A. Libby, and Xiaolei Su. The physical landscape of car-t synapse. Biophysical Journal, 123:2199–2210, 8 2024. ISSN 0006-3495. doi: 10.1016/J.BPJ.2023.09.004. URL https://doi.org/10.1016/J.BPJ.2023.09.004.

[23] Z. Wang, R. Lu, W. Wang, F. B. Tian, J. J. Feng, and Y. Sui. A computational model for the transit of a cancer cell through a constricted microchannel. Biomechanics and Modeling in Mechanobiology, 22:1129–1143, 8 2023. ISSN 16177940. doi: 10.1007/S10237-023-01705-6. URL https://doi.org/10.1007/s10237-023-01705-6.

[24] S. Hervas-Raluy, J. M. Garcia-Aznar, and M. J. Gomez-Benito. Modelling actin polymerization: the effect on confined cell migration. Biomechanics and Modeling in Mechanobiology, 18:1177–1187, 8 2019. ISSN 16177940. doi: 10.1007/S10237-019-01136-2/. URL https://doi.org/10.1007/S10237-019-01136-2/.

[25] Francisco Serrano-Alcalde, José Manuel García-Aznar, and María José Gómez-Benito. The role of nuclear mechanics in cell deformation under creeping flows. Journal of Theoretical Biology, 432:25–32, 11 2017. ISSN 0022-5193. doi: 10.1016/J.JTBI.2017.07.028. URL https://doi.org/10.1016/J.JTBI.2017.07.028.

[26] Francisco Serrano-Alcalde, José Manuel García-Aznar, and María José Gómez-Benito. Cell biophysical stimuli in lobopodium formation: a computer based approach. Computer Methods in Biomechanics and Biomedical Engineering, 24:496–505, 2021. ISSN 14768259. doi: 10.1080/10255842.2020.1836622. URL https://doi.org/10.1080/10255842.2020.1836622.

[27] Thomas Rüberg and José Manuel Garcí Aznar. Numerical simulation of solid deformation driven by creeping flow using an immersed finite element method. Advanced Modeling and Simulation in Engineering Sciences, 3:1–31, 12 2016. ISSN 22137467. doi: 10.1186/S40323-016-0061-0. URL https://doi.org/10.1186/S40323-016-0061-0.

[28] Daniel Camacho-Gomez, Nieves Movilla, Carlos Borau, Alejandro Martin, Carmen Oñate Salafranca, Julian Pardo, Maria Jose Gomez-Benito, and Jose Manuel Garcia-Aznar. An agent-based method to estimate 3d cell migration trajectories from 2d measurements: Quantifying and comparing t vs car-t 3d cell migration. Computer Methods and Programs in Biomedicine, 255:108331, 10 2024. ISSN 0169-2607. doi: 10.1016/J.CMPB.2024.108331. URL https://doi.org/10.1016/J.CMPB.2024.108331.

[29] Francisco Merino-Casallo, Maria Jose Gomez-Benito, Ruben Martinez-Cantin, and Jose Manuel Garcia-Aznar. A mechanistic protrusive-based model for 3d cell migration. European Journal of Cell Biology, 101:151255, 6 2022. ISSN 0171-9335. doi: 10.1016/J.EJCB.2022.151255. URL https://doi.org/10.1016/J.EJCB.2022.151255.

[30] Daniel Camacho-Gomez, Raffaele Sentiero, Maurizio Ventre, and Jose Manuel Garcia-Aznar. Leveraging agent-based models and deep reinforcement learning to predict taxis in cell migration. npj Systems Biology and Applications 2025 11:1, 11:1–8, 8 2025. ISSN 2056-7189. doi: 10.1038/s41540-025-00576-0. URL https://doi.org/10.1038/s41540-025-00576-0.

[31] Katsuhiko Sato. A cell membrane model that reproduces cortical flow-driven cell migration and collective movement. Frontiers in Cell and Developmental Biology, 11:1126819, 6 2023. ISSN 2296634X. doi: 10.3389/FCELL.2023.1126819. URL https://doi.org/10.3389/FCELL.2023.1126819.

[32] Ismael González-Valverde and José Manuel García-Aznar. Mechanical modeling of collective cell migration: An agent-based and continuum material approach. Computer Methods in Applied Mechanics and Engineering, 337:246–262, 8 2018. ISSN 0045-7825. doi: 10.1016/J.CMA.2018.03.036. URL https://doi.org/10.1016/J.CMA.2018.03.036.

[33] Yousef Jamali, Mohammad Azimi, and Mohammad R.K. Mofrad. A sub-cellular viscoelastic model for cell population mechanics. PLOS ONE, 5:e12097, 2010. ISSN 1932-6203. doi: 10.1371/JOURNAL.PONE.0012097. URL https://doi.org/10.1371/JOURNAL.PONE.0012097.

[34] Jorge Escribano, Raimon Sunyer, María Teresa Sánchez, Xavier Trepat, Pere Roca-Cusachs, and José Manuel García-Aznar. A hybrid computational model for collective cell durotaxis. Biomechanics and Modeling in Mechanobiology, 17:1037–1052, 8 2018. ISSN 16177940. doi: 10.1007/S10237-018-1010-2. URL https://doi.org/10.1007/S10237-018-1010-2.

[35] Sandipan Chattaraj, Julius Zimmermann, and Francesco Silvio Pasqualini. Biopoint: A particle-based model for probing nuclear mechanics and cell-ecm interactions via experimentally derived parameters. bioRxiv, page 2025.04.22.650069, 4 2025. doi: 10.1101/2025.04.22.650069. URL https://doi.org/10.1101/2025.04.22.650069.

[36] Sami Alawadhi, David M. Rutkowski, Yerbol Tagay, Alexander X. Cartagena-Rivera, Alexander S. Zhovmer, Denis Tsygankov, Dimitrios Vavylonis, and Erdem D. Tabdanov. Immune cell migration models synergize nuclear piston, uropod, and microenvironment into hydraulic cell engine. bioRxiv, page 2025.09.02.673867, 9 2025. ISSN 2692-8205. doi: 10.1101/2025.09.02.673867. URL https://doi.org/10.1101/2025.09.02.673867.

[37] Oleg Mikhajlov, Ram M. Adar, Maria Tătulea-Codrean, Anne-Sophie Macé, John Manzi, Fanny Tabarin, Aude Battistella, Fahima di Federico, Jean-François Joanny, Guy Tran van Nhieu, and Patricia Bassereau. Cell adhesion and spreading on fluid membranes through microtubules-dependent mechanotransduction. Nature Communications 2025 16:1, 16:1–17, 1 2025. ISSN 2041-1723. doi: 10.1038/s41467-025-56343-6. URL https://doi.org/10.1038/s41467-025-56343-6.

[38] Patrick R. O’Neill, Jean A. Castillo-Badillo, Xenia Meshik, Vani Kalyanaraman, Krystal Melgarejo, and N. Gautam. Membrane flow drives an adhesion-independent amoeboid cell migration mode. Developmental Cell, 46:9–22.e4, 7 2018. ISSN 1534-5807. doi: 10.1016/J.DEVCEL.2018.05.029. URL https://doi.org/10.1016/J.DEVCEL.2018.05.029.

[39] Henry De Belly, Shannon Yan, Hudson Borja da Rocha, Sacha Ichbiah, Jason P. Town, Patrick J. Zager, Dorothy C. Estrada, Kirstin Meyer, Hervé Turlier Carlos Bustamante, and Orion D. Weiner. Cell protrusions and contractions generate long-range membrane tension propagation. Cell, 186:3049–3061.e15, 7 2023. ISSN 10974172. doi: 10.1016/J.CELL.2023.05.014/. URL https://doi.org/10.1016/J.CELL.2023.05.014/.

[40] Ahmadreza Ghaffarizadeh, Randy Heiland, Samuel H Friedman, Shannon M Mumenthaler, and Paul Macklin. Physicell: An open source physics-based cell simulator for 3-d multicellular systems. PLoS computational biology, 14(2):e1005991, 2018. doi: 10.1371/journal.pcbi.1005991. URL https://doi.org/10.1371/journal.pcbi.1005991.

[41] Takuya Akiba, Shotaro Sano, Toshihiko Yanase, Takeru Ohta, and Masanori Koyama. Optuna: A next-generation hyperparameter optimization framework. Proceedings of the ACM SIGKDD International Conference on Knowledge Discovery and Data Mining, pages 2623–2631, 7 2019. doi: 10.1145/3292500.3330701. URL https://doi.org/10.1145/3292500.3330701.

[42] James Bergstra, Rémi Bardenet, Yoshua Bengio, and Balázs Kégl. Algorithms for hyper-parameter optimization. Advances in Neural Information Processing Systems, 24, 2011. URL https://proceedings.neurips.cc/paper_files/paper/2011/file/86e8f7ab32cfd12577bc2619bc635690-Paper.pdf.

[43] Silvia Hervas-Raluy, Barbara Wirthl, Pedro E. Guerrero, Gil Robalo Rei, Jonas Nitzler, Esther Coronado, Jaime Font de Mora Sainz, Bernhard A. Schrefler, Maria Jose Gomez-Benito, Jose Manuel Garcia-Aznar, and Wolfgang A. Wall. Tumour growth: An approach to calibrate parameters of a multiphase porous media model based on in vitro observations of neuroblastoma spheroid growth in a hydrogel microenvironment. Computers in Biology and Medicine, 159:106895, 6 2023. ISSN 0010-4825. doi: 10.1016/J.COMPBIOMED.2023.106895. URL https://doi.org/10.1016/J.COMPBIOMED.2023.106895.

[44] Francisco Merino-Casallo, Maria J. Gomez-Benito, Yago Juste-Lanas, Ruben Martinez-Cantin, and Jose M. Garcia-Aznar. Integration of in vitroand in silicomodels using bayesian optimization with an application to stochastic modeling of mesenchymal 3d cell migration. Frontiers in Physiology, 9:369287, 9 2018. ISSN 1664042X. doi: 10.3389/FPHYS.2018.01246. URL https://doi.org/10.3389/FPHYS.2018.01246.

[45] Yoshihiko Ozaki, Yuki Tanigaki, Shuhei Watanabe, Masahiro Nomura, and Masaki Onishi. Multiobjective tree-structured parzen estimator. Journal of Artificial Intelligence Research, 73:1209–1250, 4 2022. ISSN 1076-9757. doi: 10.1613/JAIR.1.13188. URL https://doi.org/10.1613/JAIR.1.13188.

[46] Matthew R. Zanotelli, Aniqua Rahman-Zaman, Jacob A. VanderBurgh, Paul V. Taufalele, Aadhar Jain, David Erickson, Francois Bordeleau, and Cynthia A. Reinhart-King. Energetic costs regulated by cell mechanics and confinement are predictive of migration path during decision-making. Nature Communications, 10:4185–, 12 2019. ISSN 20411723. doi: 10.1038/S41467-019-12155-Z;TECHMETA. URL https://doi.org/10.1038/S41467-019-12155-Z;TECHMETA.

[47] Yuehua Yang and Hongyuan Jiang. Mechanical properties of external confinement modulate the rounding dynamics of cells. Biophysical Journal, 120:2306–2316, 6 2021. ISSN 15420086. doi: 10.1016/j.bpj.2021.04.006. URL https://doi.org/10.1016/j.bpj.2021.04.006.

[48] Chieh Ren Hsia, Jawuanna McAllister, Ovais Hasan, Julius Judd, Seoyeon Lee, Richa Agrawal, Chao Yuan Chang, Paul Soloway, and Jan Lammerding. Confined migration induces heterochromatin formation and alters chromatin accessibility. iScience, 25:104978, 9 2022. ISSN 2589-0042. doi: 10.1016/J.ISCI.2022.104978. URL https://doi.org/10.1016/J.ISCI.2022.104978.

[49] Claire Leclech, Giulia Cardillo, Bettina Roellinger, Xingjian Zhang, Joni Frederick, Kamel Mamchaoui, Catherine Coirault, Abdul I Barakat, C Leclech, G Cardillo, B Roellinger, X Zhang, J Frederick, AI Barakat, K Mamchaoui, and C Coirault. Micro-scale topography triggers dynamic 3d nuclear deformations. Advanced Science, 12:2410052, 3 2025. ISSN 21983844. doi: 10.1002/ADVS.202410052. URL https://doi.org/10.1002/ADVS.202410052.

[50] Gabriela Da Silva André and Céline Labouesse. Mechanobiology of 3d cell confinement and extracellular crowding. Biophysical Reviews, 16:833, 12 2024. ISSN 18672469. doi: 10.1007/S12551-024-01244-Z. URL https://doi.org/10.1007/S12551-024-01244-Z.

